# Length-dependent RNA foci formation and RAN translation are associated with SCA12 disorder

**DOI:** 10.1101/2023.07.21.550051

**Authors:** Manish Kumar, Shweta Sahni, Vivek Anand, Deepak Kumar, Neetu Kushwah, Divya Goel, Himanshi Kapoor, Achal K Srivastava, Mohammed Faruq

**Author notes:** Correspondence to: Dr. Mohammed Faruq, Address: Genomics and Molecular Medicine, CSIR-Institute of Genomics and Integrative Biology (CSIR -IGIB), Mall Road, Delhi 110007, India.

## Abstract

Spinocerebellar ataxia type-12 (SCA-12) is a neurodegenerative disease caused by tandem CAG repeat expansion in the 5’-UTR/non-coding region of *PPP2R2B*. Molecular pathology of SCA12 has not been studied in the context of CAG repeats and no appropriate models exist. We found in human SCA12-iPSC derived neuronal lineage that expanded CAG in *PPP2R2B* transcript forms nuclear RNA foci and were found to sequester variety of proteins. Further, the ectopic expression of transcript containing varying length of CAG repeats exhibits non-canonical Repeat Associated Non-AUG (RAN) translation in multiple frames in HEK293T cells, which was further validated in patient-derived neural stem cells using specific antibodies. mRNA sequencing of the SCA12 and control neurons have shown a network of crucial transcription factors affecting neural fate, in addition to alteration of various signaling pathways involved in neurodevelopment. Altogether, this study identifies the molecular signatures of spinocerebellar ataxia type-12 disorder using patient-derived neuronal cell lines.

## Introduction

Spinocerebellar ataxia-12 (SCA12) is a progressive late-onset autosomal dominantly inherited neurodegenerative disorder characterized by progressive hand tremors, mild to moderate gait ataxia, dysarthria, bradykinesia, hyperreflexia & other neuropsychiatric symptoms [1]. Spinocerebellar ataxia type-12 is caused by a CAG repeat expansion within the 5’-UTR region of the *PPP2R2B* gene on chr5q which in healthy control is 4 to 32 repeats, whereas the pathogenic threshold for the disease onset is 43 [2]. Clinically, Spinocerebellar ataxia-12 is a unique ataxia subtype with characteristic features of action tremor of hands in addition to the cerebellar and neuronal features. SCA12 disease exhibits two broad phenotypes in patientsviz. the tremor dominant or the gait dominant. Typically, the tremor and ataxia syndrome ensues in the 4th-5th decade of the patient’s life [3]. While most cases of Spinocerebellar ataxia type-12 have been reported from the Indian subcontinent, a variety of rare presentations have been reported from different parts of the world [4].

The only study on the *post-mortem* brain of Spinocerebellar ataxia-12 patients reported Purkinje cell degeneration with lesions in the cerebellar-cortical regions [1]. The same study also reported ubiquitin-positive inclusions in pars compacta and Purkinje cells, negative for polyglutamine, tau, and transactive response DNA binding protein 43 (TDP-43) [1]. Also, it was reported that the CAG repeats act as a promoter element and can increase the expression of the *PPP2R2B* gene [1]. Another study on the plasma of Spinocerebellar ataxia type-12 suggested the altered protein abundance of transportation and lipid metabolism proteins [5]. The exact molecular pathology of CAG expansion in *PPP2R2B* is unknown, however, a gain-of-function mechanism has been hypothesized. Lin et al reported that CAG repeats in a reporter construct exhibit an increase in the expression akin to the CGG repeat expansion effect on the FMR1 locus of fragile X associated Tremor ataxia syndrome (Fragile X-Associated Tremor/Ataxia Syndrome) [6]. At the protein level, the effect of CAG repeats has not been elucidated.At the clinical level, it has similarities with Fragile X-Associated Tremor/Ataxia Syndrome as well as spinocerebellar ataxias [7], however at the molecular level no similar mechanism has been so far elucidated.

We sought to decipher the molecular pathology of CAG repeat expansion within *PPP2R2B* in Spinocerebellar ataxia type-12 using patient iPSC-derived models of neuronal lineage. The pertinent questions were i) to explore the molecular correlates of tremor-ataxia phenotype, if it has a matching molecular mechanism as reported for CGG repeats in the 5’UTR of the *FMR1* gene where an RNA gain-of-function mechanism is well established, ii) to explore if the presence of CAG repeat motif exhibits polyglutamine (polyQ) pathology with protein level toxicity and thus, phenotypic equivalence with other SCA types i.e. SCA1-3 and Huntington disease, etc. In the present study, we utilized in-vitro and iPSC derived Neural stem cell lineage models of Spinocerebellar ataxia type-12 cell lines to reveal the occurrence of RNA-mediated gain-of-function through RNA-foci and, non-canonical non-ATG mediated translation of *PPP2R2B* with peptide frame coding for CAG repeats. We were able to detect the occurrence of RNA foci formation in the nucleus of the Spinocerebellar ataxia type-12 patient-derived neural stem cell model. We further explored the pathways that might be affected in the Spinocerebellar ataxia type-12 disease biology through RNA pull-down assay and transcriptomics analysis. In addition, we depicted the process of RAN translation (Repeat Associated Non-AUG mediated translation), which has been reported in other repeat expansion disorders including Fragile X-Associated Tremor/Ataxia Syndrome [8] [9], in the *PPP2R2B* gene driven by the expanded CAG repeat motif, both in the transient overexpression model of *PPP2R2B* and in the patient-derived cellular model of Spinocerebellar ataxia type-12. In summary, our study provides preliminary evidence linking the molecular mechanisms of disease pathology to the tremor-ataxia phenotype as in Fragile X-Associated Tremor/Ataxia Syndrome and other PolyQ diseases and also provides clues towards the global molecular disruptions caused by the pathogenic CAG repeat.

## Results and Discussion

### Pathogenic Spinocerebellar ataxia type-12 repeats induce RNA foci formation in the nucleus

Repeat-containing transcripts are known to form nuclear aggregates in the form of RNA foci in many repeat-mediated CAG/CUG diseases [23]. To check for the occurrence of similar repeat-containing transcript aggregates in our Spinocerebellar ataxia type-12 NSC model, we designed probes against the CAG-repeat region. We utilized the RNA FISH technique to determine the location of CAG-containing transcript within the NSCs generated from three patients and two controls with varying repeat lengths (Table 1). Spinocerebellar ataxia type-12 positive cell lines showed the presence of RNA foci or aggregate-like formations inside the nucleus of cells (Figure 1A, 1B, 1C). These aggregates were lacking in the secondary neuronal cell lines (SK-N-SH) (Figure 1D) and control cell lines and (Figure 1E and 1F).

**Table 1.** Sample details and associated metadata.

| **Sample ID** | **Cell line ID** | **CAG Repeat** | **Design** | **Condition** | **Cell_line** | **Sex** | **Batch** |
| --- | --- | --- | --- | --- | --- | --- | --- |
| 10223NSC-1 | 10223 NSC | 09/16 | Control_NSC | Control | NSC | MALE | Batch 2 |
| 10223NSC-2 | 10223 NSC | 09/16 | Control_NSC | Control | NSC | MALE | Batch 2 |
| 21-14NSC-2 | 21/14 NSC | 17/65 | Diseased_NSC | Diseased | NSC | MALE | Batch 2 |
| 21-14NSC-1 | 21/14 NSC | 17/65 | Diseased_NSC | Diseased | NSC | MALE | Batch 2 |
| 2555NEU | 2555 Neuron | 14/59 | Diseased_Neuron | Diseased | Neuron | MALE | Batch 2 |
| 2555NSC-1 | 2555 NSC | 14/59 | Diseased_NSC | Diseased | NSC | MALE | Batch 2 |
| 2555NSC-2 | 2555 NSC | 14/59 | Diseased_NSC | Diseased | NSC | MALE | Batch 2 |
| D161NSC-1 | D161 NSC | 10/13 | Control_NSC | Control | NSC | FEMALE | Batch 2 |
| D161NSC-2 | D161 NSC | 10/13 | Control_NSC | Control | NSC | FEMALE | Batch 2 |
| SRR5754278 | 10223 iPSC | 09/16 | Control_iPSC | Control | iPSC | MALE | Batch 1 |
| SRR5754279 | 10223 iPSC | 09/16 | Control_iPSC | Control | iPSC | MALE | Batch 1 |
| SRR5754280 | 10223 NSC | 09/16 | Control_NSC | Control | NSC | MALE | Batch 1 |
| SRR5754281 | 10223 NSC | 09/16 | Control_NSC | Control | NSC | MALE | Batch 1 |
| SRR5754282 | 10223 Neuron | 09/16 | Control_Neuron | Control | Neuron | MALE | Batch 1 |
| SRR5754283 | 10223 Neuron | 09/16 | Control_Neuron | Control | Neuron | MALE | Batch 1 |
| SRR5754576 | 2555 iPSC | 14/59 | Diseased_iPSC | Diseased | iPSC | MALE | Batch 1 |
| SRR5754577 | 2555 iPSC | 14/59 | Diseased_iPSC | Diseased | iPSC | MALE | Batch 1 |
| SRR5754626 | 2555 NSC | 14/59 | Diseased_NSC | Diseased | NSC | MALE | Batch 1 |
| SRR5754627 | 2555 Neuron | 14/59 | Diseased_Neuron | Diseased | Neuron | MALE | Batch 1 |
| SRR57546<br>28 | 2555 Neuron | 14/59 | Diseased_Neuron | Diseased | Neuron | MALE | Batch<br>1 |
| SRR57547<br>16 | 11/08 iPSC | 10/67 | Diseased_iPSC | Diseased | iPSC | FEMALE | Batch<br>1 |
| SRR57547<br>17 | 11/08 iPSC | 10/67 | Diseased_iPSC | Diseased | iPSC | FEMALE | Batch<br>1 |
| SRR57547<br>18 | 21/14 iPSC | 17/65 | Diseased_iPSC | Diseased | iPSC | MALE | Batch<br>1 |

**Figure 1.**
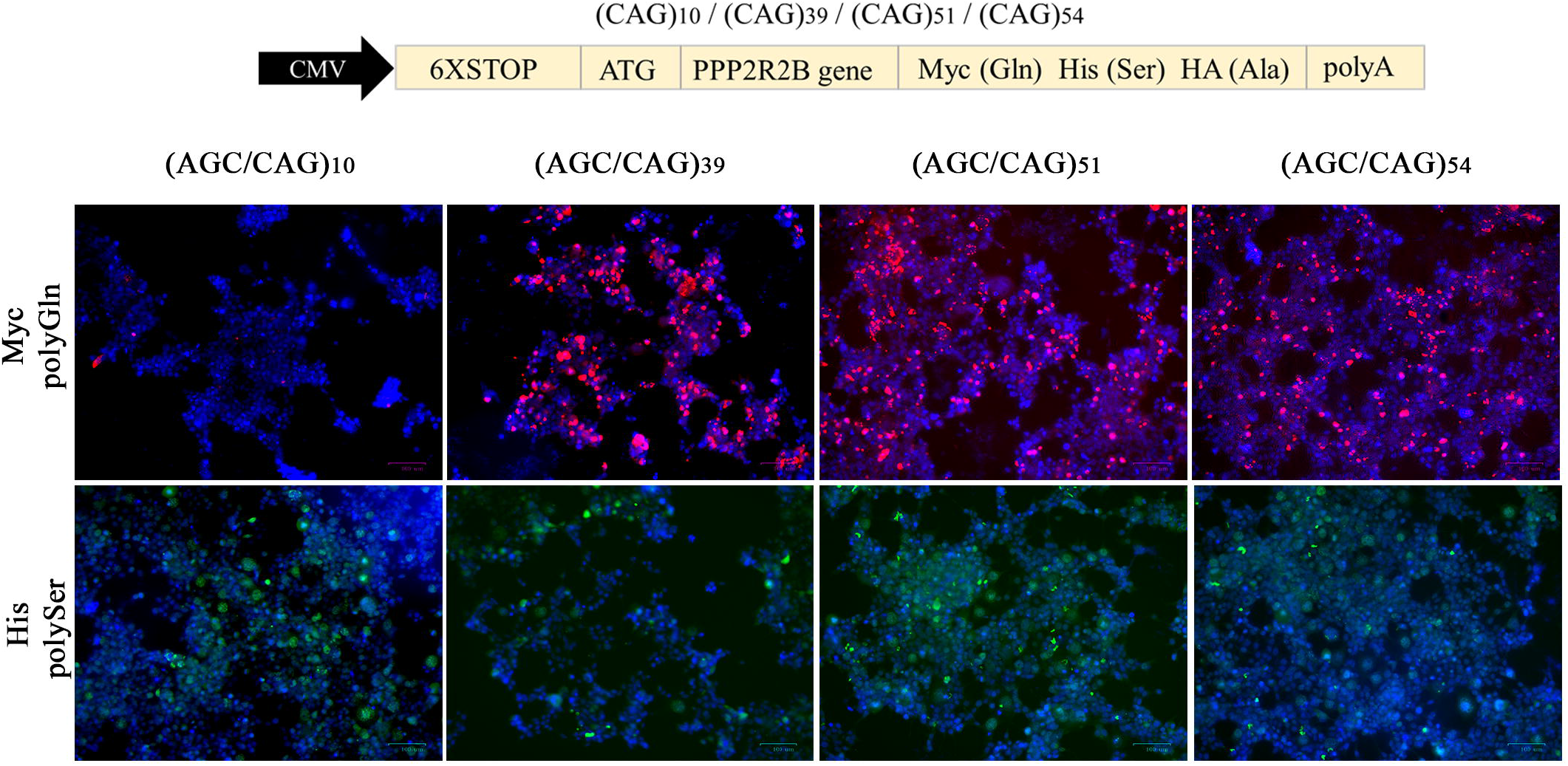
RNA FISH analysis of SCA12 Positive Neural Stem Cells shows RNA aggregates inside the nucleus of the cells. Where **(A)(B)** and **(C)** are SCA12 Positive Neural Stem cell lines with CAG(14/59), CAG (10/67) and CAG (17/65) respectively. **(D)** SK-N-SH cell line. **(E)** and **(F)** are two healthy control derived NSCs with CAG (09/16) and CAG (10/13) respectively. CTG probe labeled with CY3 shown in Red, and DAPI is shown in Blue with magnification bar of 25 µM (micrometer).All of the images exhibited here have been converted as MIP (maximum intensity projection) images for enhanced visualization using Nikon’s NIS elements software.

RNA foci are considered a hallmark of RNA-mediated toxicity in other neurodegenerative diseases, their presence in the Spinocerebellar ataxia type-12 disease model links it to the disease pathogenesis. Further, we checked the colocalization of MBNL1(muscleblind-like 1) protein, the protein most commonly known to be associated with RNA foci in other repeat-mediated diseases [23] with the *PPP2R2B* RNA foci. RNA FISH combined with IF for MBNL1 protein using anti-MBNL1 antibody did not find colocalization of MBNL1 with SCA-12 RNA foci (Mander’s overlap equation 0.5) (Data not shown).

### Expanded Spinocerebellar ataxia type-12 CAG repeats induced RNA foci to bind key nuclear proteins

To determine if the Spinocerebellar ataxia type-12 nuclear RNA foci in NSCs interact with or bind any specific proteins within the cellular milieu, we performed RNA-pull down assay. CAG repeats with their flanking regions upstream and downstream of the *PPP2R2B* gene from the patient (CAG-54) and control DNA (CAG-10) was cloned into pcDNA3.1 vector and amplified using primers against T7 promoter and PPP2R2B region. Next, this DNA template was used to make RNA through the Hi-scribe transcription kit using biotinylated-CTP (Thermo Fisher) instead of dCTP. Both the RNAs (CAG-54 and CAG-10) were incubated with nuclear proteins extracted from Spinocerebellar ataxia type-12 positive NSCs and control NSCs, and a pull-down was performed using streptavidin beads to check for interaction through mass spectrometry. The high score peptides obtained from the mass spectrometry-based detection of interacting proteins revealed that 378 proteins from control-NSC nuclear protein extract were bound in high concentration with the CAG-10 construct, (CT_10= 378) while 623 proteins from the Spinocerebellar ataxia type-12 positive NSC protein extract were retained by the CAG-10 construct (SCA12_10=623). Of the total interacting proteins observed, 303 proteins were common in both the groups that were found to bind to the CAG-10 repeat (CT_10 ∩ SCA12_10= 303). Similarly, we looked for the identity and number of proteins retained by the pathogenic CAG repeat construct (CAG-54). 362 proteins were found to be associated with CAG-54 repeats from the nuclear extract of control NSC in high concentration (CT_54= 362). 179 proteins from Spinocerebellar ataxia type-12 positive NSC nuclear protein extract were found to bind with CAG-54 repeats (SCA12_54= 179), of which 115 proteins were common in both the groups (CT_54 ∩ SCA12_54 = 115). Further analysis revealed that 13 proteins exclusively bind to CAG-54 repeats (KRT27, HSPA1B, H1-3, DYNLL2, C1QBP, S100A16, CTSA, KIAA0100, PSMA2, FN1, SNRPB, GSN, and PSMA1) (Supplementary Figure 5).

Further investigation of these proteins revealed that their accumulation had previously been associated with several neurodegenerative disorders such as Parkinson’s disease, Alzheimer’s disease, and ALS. Hsp70 proteins, particularly HSPA1B, are important in protein folding and preventing misfolded proteins from aggregating. These proteins aid in the refolding of damaged proteins and direct their degradation by the proteasome. HSPA1B is overexpressed in the substantia nigra of patients with Parkinson’s disease and frontotemporal dementia with Parkinsonism. It has also been proposed that Hsp70 associates with protein aggregates as an endogenous effort to alleviate the toxic effects of the aggregates in Alzheimer’s and Parkinson’s brains [24]. Similarly, C1QBP expression was found to be considerably lower in Alzheimer’s in microarray analysis, despite being elevated in normal aging. C1QBP has been demonstrated to interact with Aβ and stimulate fibrillogenesis, the process by which Aβ aggregates into fibrils. This interaction indicates that C1QBP may be involved in the production and buildup of amyloid plaques in Alzheimer’s disease[25]. Furthermore, Proline-Arginine dipeptide repeat (PR-DPR), the most toxic protein formed by RAN translation in Amyotrophic Lateral Sclerosis patients with C9ORF72 gene expansion mutation is known to interact with C1QBP molecule to activate intracellular NLRP3 (NOD-, LRR-, and pyrin domain-containing 3) inflammasome activation of microglia cells [26]. DYNLL2 (Dynein light chain LC3 type-2) is a motor protein involved in vesicle and organelle axonal-retrograde transport. Many studies have linked dynein to motor neuron disease, Parkinson’s disease, and Alzheimer’s disease. To begin, interruption of axonal transport is a common hallmark of these neurodegenerative illnesses. Second, autophagy and the clearance of aggregation-prone proteins, both of which rely on dynein proteins, are reported to be defective in these disorders [27]. The FN1 (Fibronectin 1) molecule is a glycoprotein found in the extracellular matrix that is involved in cell adhesion, tissue organization, and migration. It participates in clotting, which may influence amyloid-beta fibrillization [28]. Furthermore, FN1 expression was shown to be higher in the plasma of Alzeimer’s patients with dementia compared to Alzeimer’s patients without dementia symptoms [28].In addition, FN1 has been linked to motor neuron degeneration after being activated by the pro-inflammatory TGF-β pathway [29].

SNRPB (Small Nuclear Ribonucleoprotein Polypeptides B), on the other hand, is involved in the assembly and functioning of the spliceosome complex. Unavailability or dysfunction of this protein may result in abnormal splicing processes, leading to hyperexcitability and cognitive impairment in Alzheimer’s disease [30]. GSN (Gelsolin) is an actin-binding protein that plays a role in cytoskeleton dynamics, cell motility, and cellular responses to extracellular signals. GSN is known to inhibit Amyloid aggregation and cellular apoptosis in Alzheimer’s disease. Also, GSN is observed to have reduced expression in advanced disease states of AD leading to exacerbation of disease [31]. Similarly, S100A6 (Calcyclin or Calmodulin-like protein 3) is a member of the S100 family of calcium-binding proteins whose expression level has been observed to be altered in Alzheimer’s Disease, Parkinson’s Disease, Amyotrophic Lateral Sclerosis, and Huntington’s disease. Recently, its role has been discovered in the differentiation of astrocytes [32].Furthermore, it has recently been discovered that RNA foci emerge from the nucleus to cytoplasm over time and that RNA and RAN protein interaction and aggregation are associated with RNA binding protein dispositioning and cellular toxicity[33].This shows that RNA and RAN proteins interact to cause and aggravate disease over time.

Overall, these are critical proteins with important functions to perform in the cell, and their absence or altered expression can cause serious disruptions in cellular processes, as shown in other neurodegenerative disorders.

### RAN Proteins Accumulate in Neural stem Cells with expanded CAG repeats

To check if the expanded *PPP2R2B* leads to RAN translation in transfected cells, multiple constructs with different CAG/AGC repeats sizes of *PPP2R2B* mini genes (60, 50, 43, and 10) were generated with 3 epitope tags in each reading frame (Supplementary 1) (Figure 2A) and characterized using the Sanger sequencing method (Data not shown). As reported previously, CAG repeat length decreases when amplified inside a plasmid using bacterial culture [34]. We also observed much heterogeneity in the sequences of picked colonies after transformation, hence we proceeded with CAG repeat lengths of 54, 51, 39, and 10 repeats. According to the NCBI and UniProt databases, the CAG repeat (Glutamine) falls under the reading frame of AGC (Serine), hence we have referred to the CAG repeat sequence as the AGC repeat sequence in Figure 2B. Transient overexpression of RAN constructs in the HEK293T cell line revealed the occurrence of RAN translation within the CAG/AGC extended reading frame which emit positive signals unique to polyGlu and polySer proteins. Surprisingly, polyGlu and polySer proteins were expressed in constructs lacking the ATG start codon in the enlarged CAG/AGC repeats. Also, PolyGlutamine was found to be expressed when the only start codon ATG was mutated to GGG, this suggests that RAN Translation of the PolyGlutamine frame is favored in the *PPP2R2B* gene inside the HEK293T cell line (Supplementary Figure 6). CAG/AGC with unexpanded repeats (10), on the other hand, did not create any RAN proteins. We did not see any signal with the HA tag antibodies because the *PPP2R2B* gene sequence is predicted to induce a stop codon in the Alanine frame.

**Figure 2.**
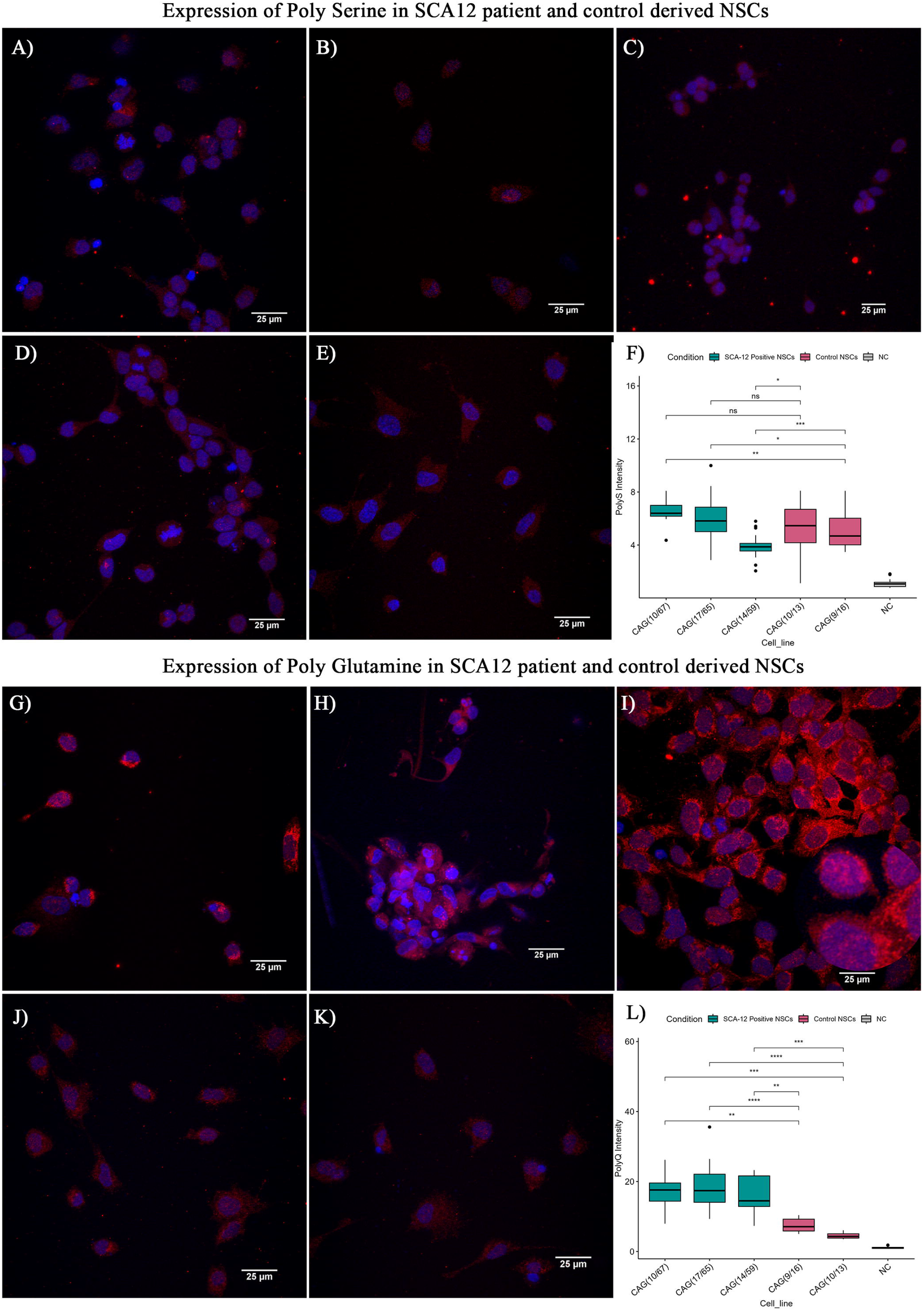
Repeat dependent RAN protein expression in HEK293T cell line. **(A)** RAN construct having 6x stop codon in 5 prime region, variable CAG repeat length followed by three different epitopes respective for each reading frame. **(B)** Immunofluorescence microscopy results of HEK293T cells after transfection with indicated constructs, where DAPI is shown in Blue, Myc/polyGln (polyglutamine) signal is shown in red and His/polySer (polyserine) is shown in green with magnification bar of 100 µM (micrometer).

To confirm the occurrence of RAN translation and its associated proteins in Spinocerebellar ataxia type-12 disease, we designed polyclonal antibodies against the anticipated C-terminal portions of the PPP2R2B gene that were specific to polySer (AGC) and polyGlu (CAG) proteins (Supplementary Figure 4) that were commercially obtained to evaluate the presence of RAN proteins in our disease model i.e. Spinocerebellar ataxia type-12 positive NSCs. These antibodies were used to perform immunocytochemistry (ICC) and confocal microscopy on Spinocerebellar ataxia type-12 NSCs and control NSCs (Figure 3). Confocal microscopy analysis showed significant difference in both PolySer and PolyGlu expression (Figure 3A-E and Figure 3G-K). PolySer intensity profile showed little higher expression of PolySer in two SCA12 NSCs (CAG 10/67 and 17/65) with less significance whereas significantly lower expression in one SCA12 NSC (14/59) which suggests repeat number variation may also play role in RAN Proteins expressions (Figure 3F). However, PolyGlu intensity profile showed a strikingly significant difference in the intensity of signals for poly-Glu RAN specific antibodies in Spinocerebellar ataxia type-12 NSCs, which was comparatively trivial in control NSCs (Figure 3L).Our data provide evidence for RAN-mediated translation of the *PPP2R2B* gene in a repeat length-dependent non-canonical manner, resulting in the synthesis of enlarged homopolymer proteins in multiple frames which might be responsible for cellular toxicity as observed in other repeat-containing diseases such as Fragile X-Associated Tremor/Ataxia Syndrome, Huntington’s disease, etc.

**Figure 3.**
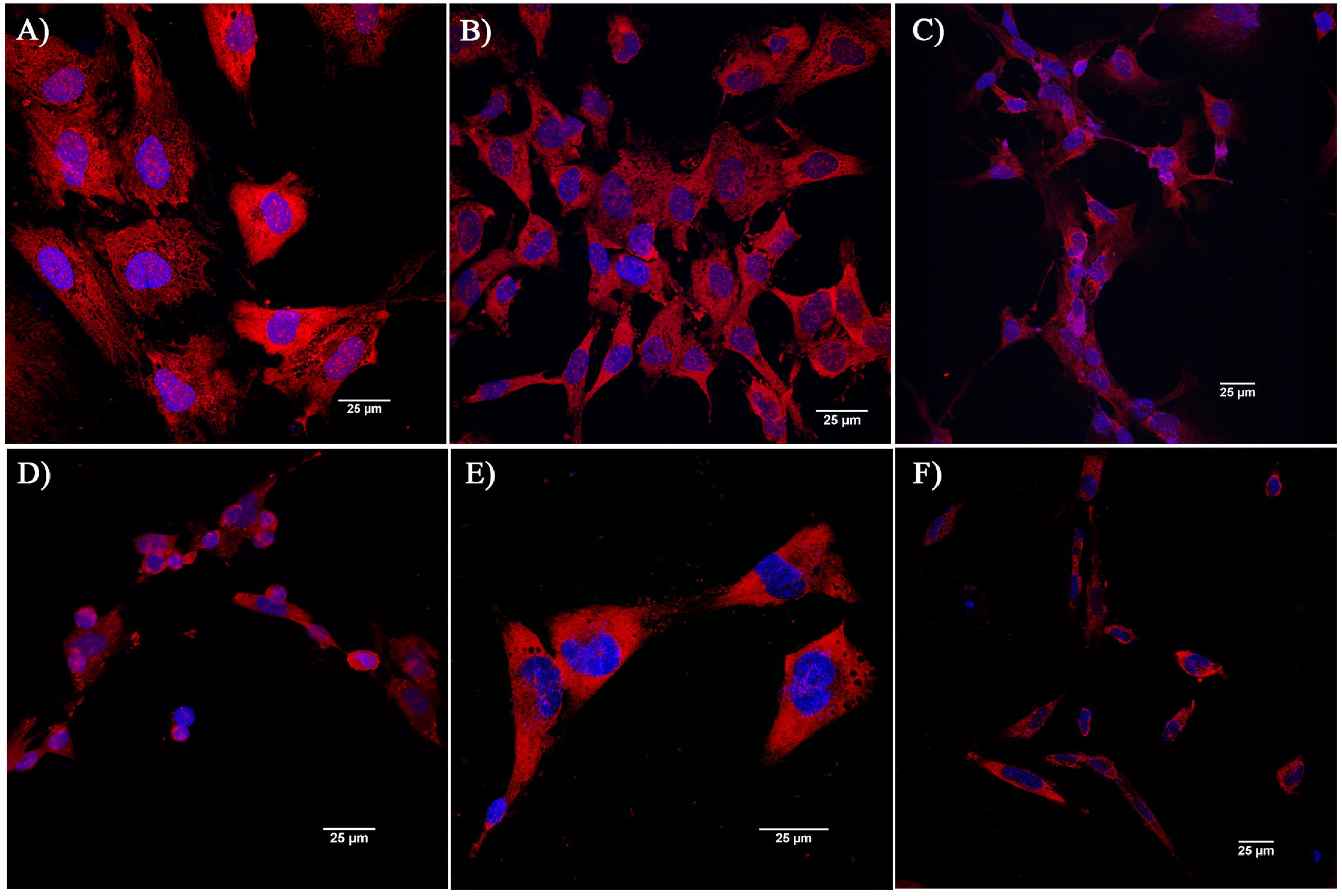
Expression of RAN proteins in SCA12 vs Control Neuronal cell lines. Where **(A)** to **(E)** are cell lines showing positive staining for red (PolySerine) and blue (DAPI). **(A)(B)** and **(C)** are SCA12 Positive Neural Stem cell lines with CAG(14/59), CAG (10/67) and CAG (17/65) respectively. **(D)** and **(E)** are two healthy control derived NSCs with CAG (09/16) and CAG (10/13) respectively. **(F)** This graph compares the intensity of PolyS (Poly Serine) in Case and Control cell lines using student t-test (* represents p-value < 0.05) **(G)** to **(E)** are cell lines showing positive staining for red (Polyglutamine) and blue (DAPI) **(G)(H)** and **(I)** are SCA12 Positive Neural Stem cell lines with CAG(14/59), CAG (10/67) and CAG (17/65) respectively. **(J)** and **(K)** are two healthy control derived NSCs with CAG (09/16) and CAG (10/13) respectively. All of the images exhibited here have been converted as MIP (maximum intensity projection) images for enhanced visualization using Nikon’s NIS elements software by Nikon with magnification bar of 25 µM (micrometer) **(L)** This graph compares the intensity of PolyQ (Poly Glutamine) in Case and Control cell lines using student t-test. (* represents p-value < 0.05)

### Expression of expanded CAG repeats induces cell death

Annexin-V mediated apoptosis assay was used to determine if RAN proteins induce apoptosis in cells showing RAN translation. We transfected the HEK293T cell line with multiple RAN constructs of different CAG lengths (54, 51, 32, 10), after 36 hours, cells were treated with staurosporine for 6 hours. Cells were then trypsinized and 5ul of Annexin V with 20ul of binding buffer was added and then the percentage of apoptotic cells was detected using BD-Accuri C6 plus FACS (Fluorescence-activated cell sorting) (Supplementary Figure 7), the percentage of apoptotic cells was significantly higher in expanded CAG construct (54, 51) in comparison to the normal repeat length of CAG (10). This indicates that a mutated PPP2R2B mini gene with a CAG length of > 51 upon expression through RAN translation potentially induces cellular death in HEK293T cell lines.

### Widespread transcriptional alterations are present in Spinocerebellar ataxia type-12 derived pan neurons and neuronal progenitors

To study the changes in global transcriptional signatures induced by the pathogenic CAG repeats, mRNA sequencing was performed. The sample set consisted of four, four, and two biological replicates for iPSC, NSC, and neuronal lineage respectively along with their technical replicates, details appended in Table-1. RNA isolated from all the samples was subjected to PolyA enrichment-based mRNA sequencing. An average of 38 million sequencing reads were generated for all samples taken together and approximately 83% of the reads were mapped to the human genome (Table-1). After processing the sequencing data to reduce noise, remove outliers and normalize, batch effect correction was performed using Limma to remove the variation introduced due to sample collection and sequencing in two batches. Principal Component Analysis (PCA) showed a separation of patient and control samples along the PC2 (Y-axis) based on the expression of the 500 most variable genes (Supplementary Figure-8). We next evaluated the RNA-seq data in pairwise comparisons for each cell lineage using DESeq2 [35]. A total number of 20525, 20283, and 22150 genes were detected in iPSC, NSC, and Neurons respectively. Between developmental stages, the number of differentially expressed genes between patients and controls was the highest for iPSCs and the least for Neurons (Figure-4). Patient neurons and iPSCs showed more genes to be significantly upregulated, while patient NSCs showed greater downregulation of genes in comparison to controls (Figure-4). Unsupervised hierarchical clustering of the samples demonstrated a clear separation between patient and control derived cell-types (Supplementary Figure-8 B).To further query the nature of global transcriptome alterations induced by the pathogenic Spinocerebellar ataxia type-12 CAG repeats in the *PPP2R2B* gene, we performed pathway analysis using Qiagen’s Ingenuity Pathway Analysis (IPA) software. A comprehensive list of the top 30 significant pathways for neurons, NSC, and iPSC datasets are listed in Supplementary Table 2. In the neuronal dataset, IPA identified Signaling pathways as the most significant canonical pathways. Neurovascular Coupling signaling emerged as the top most significant pathway predicted to be upregulated in SCA-12 neurons, which has previously been linked to Alzheimer’s disease [36, 37]. The other top significant pathways in Spinocerebellar ataxia type-12 neurons include the GABA receptor signaling pathway, WNT/β-catenin Signaling, and CREB signaling in neurons. IPA results also highlight the involvement of pathways of immune response i.e. Neuroinflammation signaling pathway and the Antigen presentation pathway to be altered in the Spinocerebellar ataxia type-12 neurons. Neuroinflammation has been described in many neurodegenerative diseases earlier including Alzheimer’s disease, Parkinson’s disease, and Amyotrophic Lateral Sclerosis [38]. Further, Spinocerebellar ataxia type-12 neurons showed dysregulation of the SNARE signaling pathway. SNAREs have long been associated with critical neuronal functions such as synaptic transmission, neurite initiation and outgrowth, axon specification, and synaptogenesis (Supplementary Table 2) [39] [40].

IPA of the NSC dataset revealed several pathways involved in the transition of NSC to the neuronal lineage. CREB signaling in neurons, Neurovascular coupling signaling pathway, and WNT/β-catenin Signaling pathways which were identified in the neuronal dataset also appeared in the NSC top pathways by significance. Other top significant pathways identified in the NSC dataset include Synaptogenesis signaling, Endocannabinoid neuronal synapse, synaptic long-term depression, serotonin receptor signaling, axonal guidance signaling, and gap junction signaling. This is indicative of the role of pathogenic repeats in affecting the development of neuronal junctions and synapses. NSC pathways also exhibit signatures indicative of other neurodegenerative diseases such as Amyotrophic Lateral Sclerosissignaling and Multiple Sclerosis signaling pathways. NSCs showed a predicted inhibition of pathways involved in inflammatory responses such as HMGB1 signaling, IL-17 Signaling, Pathogen Induced Cytokine Storm Signaling Pathway, and Wound Healing Signaling Pathway. Necessitated by the presence of pathogenic repeats and their associated pathology i.e. RNA foci and RAN-translated proteins, inhibition of the inflammatory immune responses seems to be crucial to allow the development of NSCs into neurons.

### Different cell lineages exhibit differential PPP2R2B transcript isoform profiles under the Spinocerebellar ataxia type-12 disease context

The *PPP2R2B* gene is known to exhibit alternative splicing and various repeat-containing and non-repeat-containing variants have been reported. *PPP2R2B* isoforms differ in their 5’ sequence and therefore code for proteins with variable N-terminal regions, thus affecting their subcellular localization and substrate specificity of the PP2A enzyme [41] [42] We utilized the RNA-seq data to study the variation in transcript isoform profile of *PPP2R2B* gene across different cell-lineages induced by the presence of pathogenic CAG repeats. Mean-of-inferential replicates were used to quantify the expression difference in fold change between Spinocerebellar ataxia type-12 and control cell lines (Figure 4). The transcripts which were statistically different between patients and controls are NM_181674.2 (Variant 2), NM_181677.2 (Variant 5), and NR_073527.1 (Variant 12). NM_181674.2 (Variant 2) and NM_181677.2 (Variant 5) are both coding transcripts and their expression was reduced in the SCA-12-derived pan neurons. On the other hand, NR_073527.1 (Variant 12) is a non-coding transcript of the *PPP2R2B* gene but its expression was found to be increased in Spinocerebellar ataxia type-12 neurons. Overall, in the Spinocerebellar ataxia type-12 neurons, most isoforms of the *PPP2R2B* gene showed a downregulation, except NM_001271900.2 (Variant 8) and NR_073527.1 (Variant 12) in comparison to controls. While NM_001271900.2 (Variant 8) is a non-coding transcript of 10917 bp, its expression difference did not reach statistical significance. In contrast to the Spinocerebellar ataxia type-12 neurons, transcript profiles in Spinocerebellar ataxia type-12 NSCs showed a striking difference in terms of more than 2.5-fold upregulation of two isoforms viz. NM_181674.2 (Variant 2) and NM_181676.2 (Variant 4). Interestingly, both these are coding transcript isoforms and show a statistically significant upregulation in the patient NSCs when compared to controls. The other transcript isoforms identified in the NSCs did not show any significant change between patients and controls. We also looked at the isoform profile at the iPSC stage and found two isoforms NM_001271948.1 (Variant 10) and NR_073527.1 (Variant 12) to be strongly upregulated (L2Fc >4) in the SCA-12 iPSCs. NR_073527.1 (Variant 12) is a non-coding isoform of *PPP2R2B* but NM_001271948.1 (Variant 10) is a coding mRNA of 2578 bp. Despite limitations on the number of samples, the transcript isoform level data from our study gives interesting insights into consistent downregulation of *PPP2R2B* gene isoforms in the mature SCA-12 neurons, while many transcript isoforms show abundant expression at the progenitor stages.

**Figure 4.**
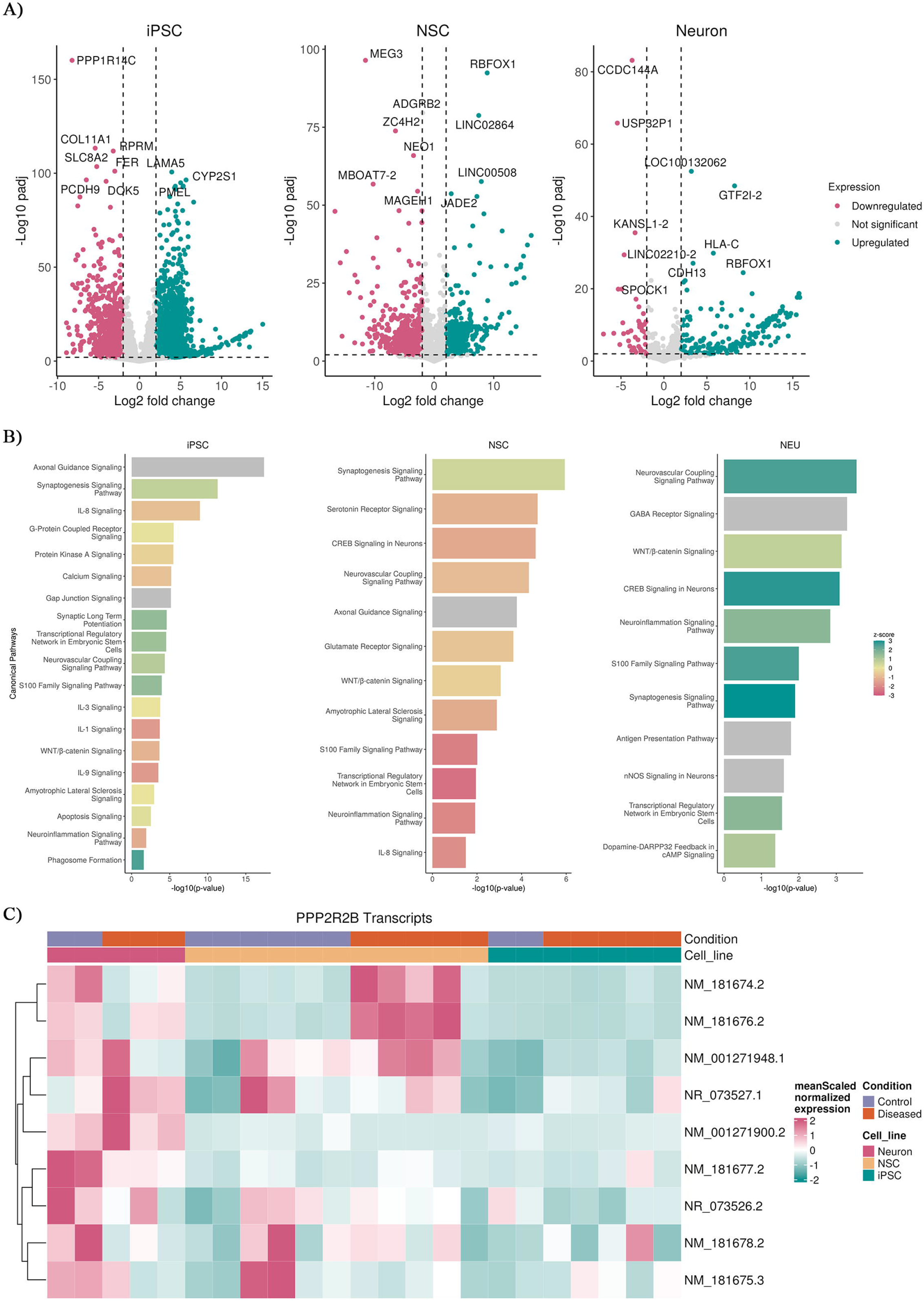
Differential expression and enrichment analysis of iPSCs, NSCs and Neurons in SCA12 disorder. **A)** Volcano plots of the differentially expressed genes within the cut off of | log2Fold change| > 2 and padj< 0.01 in iPSC, NSC and Neuron. Labeled the top 10 genes based on padj values. **B)** Barplot of enrichment of canonical pathways identified from IPA. X axis represents -log10(pvalue) and colours scaled with z-score, representing enrichment (z-score > 0) or depletion (z score < 0) of the pathway. Grey colour represents not enough genes present to calculate z-score or the direction of enrichment. **(C)**Heatmap of the PPP2R2B transcript expression in iPSC, NSC and Neuron. Values in the heatmap are scaled Mean counts of the inferential replicates.

### Transcriptional dysregulation in Spinocerebellar ataxia type-12 involves a network of Transcription Factor genes responsible for nervous system development and function

The list of DEGs was used to infer biologically and disease-relevant networks among represented genes based on their known interactions using the IPA. One of the top networks identified by the IPA is defined by the terms Nervous system development and function, organ morphology, and tissue morphology and has a top score of 42 (Supplementary Figure 9). This network has a total of 35 molecules with 26 of them picked up from our DEG gene list. Interestingly, *PPP2R2B* emerges as one of the hub genes in the network where most other genes are involved in functions related to the development, differentiation, and morphology of the nervous system and neurons. The network contains 8 genes for transcription factors viz. *BMAL1*, *GBX2*, *HIF3A*, *MYT1*, *NEUROD4*, *NEUROG2*, *PAX6* and *SOX2*. NEUROG2is a transcription factor that induces the differentiation of progenitor cells into terminal neuronal cells [43]. This lineage commitment is driven by sequential transcriptional activation of *PAX6* followed by *NEUROG2*. NEUROG2 forms a heterodimer with NEUROD4, which regulates neurogenesis in the cortex by accelerating specific transcriptional patterns [44]. Both NEUROG2 and NEUROD4 are predominantly expressed in progenitor cells. *MYT1* is a zinc-family pan-neuron-specific transcription factor that safeguards neuronal identity by repressing non-neuronal cell fates [45]. While SOX2 is essentially recognized for its role in maintaining stemness or pluripotency of somatic cells [46], recent studies have revealed its fundamental role in differentiated neurons and glia, such as the development of proper neuronal connectivity [47], axon growth, and motor neuron maturation [48]. Despite their primary role in neural progenitor cells, the overexpression of most of these TFs in mature Spinocerebellar ataxia type-12 neurons raises the question of their role in differentiated neuronal cells.

Interestingly, *BMAL1* is considered an irreplaceable clock gene that regulates circadian rhythm and rhythmic behaviors in mammals [49] [50]. However, recent studies in preclinical models indicate the role of disruption of the *BMAL1* gene to be associated with cerebellar damage and ataxia [51][52]. The role of the *BMAL1* gene is being currently investigated in neurodegenerative pathologies such as Parkinson’s syndrome and Alzheimer’s disease [53]. Furthermore, defects in the *BMAL1* gene are associated with the regulation of inflammatory and immune responses and the regulation of neuroinflammation. Disruption of the *BMAL1-* regulated dopamine signaling and its consequent impact on neuroinflammation in the brain has been proposed as a causative factor for the onset of Parkinson’s disease [54]. A possible role of the *BMAL1* gene in neuronal function could be the synchronization of the rhythmic control of transcription of other genes controlling neuronal activity and function [55]. The downregulation of*the BMAL1* gene in Spinocerebellar ataxia type-12 neurons as observed in our data provides evidence for the above hypothesis and warrants further investigation. Future studies on some of these candidate genes might be able to provide better insights into the pathogenesis of neurodegeneration in general, and Spinocerebellar ataxia type-12 associated pathology in particular.

## Conclusion

We were able to generate disease models of the Spinocerebellar ataxia type-12 disorder using human iPSC-derived cell lines in the form of repeat-containing NSCs and Neurons. Our extensive analysis using these disease models has shed new insights into the pathological mechanism of Spinocerebellar ataxia type-12 at the molecular level. The occurrence of RNA foci and RAN translation and its associated proteins are two phenomena known to be integral to the pathology of many repeat-containing diseases. Their presence in Spinocerebellar ataxia type-12 models provides an inkling of the underlying mechanisms which result in the similitude of clinical symptoms in Spinocerebellar ataxia type-12, Fragile X-Associated Tremor/Ataxia Syndrome and Huntington’s Disease. Recently, Das MR and colleagues showed that RAN translation occurs after extended retention of nuclear foci. and after localization of RNA foci in cytoplasm it interacts with RAN products to form aggregation-like molecules [33]. Our Protein pull-down data showed the entrapment of proteins that are involved in the clearance mechanism of aggregated proteins (HSPA1B, PSMA1, PSMA2).Going forth, evaluation of the transcriptome in Spinocerebellar ataxia type-12 neurons and NSCs highlights the nature of global transcriptional alterations brought about by the dysregulation of important transcription factors. Also, pathogenic phenomena in Spinocerebellar ataxia type-12 including the occurrence of RNA foci aggregates, RAN translation products, and its associated gene expression changes possibly induce inflammatory immune cascades in these cells providing a link to the neuroinflammation associated with neurodegeneration, as evident from the transcriptomic data. In concordance with this, C1QBP which activates inflammasomes in the microglia has also been found to bind specifically to the expanded CAG transcript. Further, our data provide new evidence for the linkage of *BMAL1* gene dysregulation with cerebellar dysfunction and ataxia. Overlapping results from Protein pull-down assay and RNA seq analysis in our data suggest new molecular insights into the pathogenesis of Spinocerebellar ataxia type-12 disorder. These results are, however, of preliminary nature and need to be explored further in future studies in sync with other CAG/CTG repeat expansion diseases.

## Acknowledgments

We are sincerely thankful to all the patients and their families for their participation and cooperation in this study. We are also thankful to Dr. Himanshi Kapoor for helping with Confocal microscopy, and NCBS for providing us the one control iPSC D161 (ADBSi001-A) for our experiments. We acknowledge the efforts of Mr. BharathramUppili for helping with figure compilations and Shreya Bari for helping with data analysis.

## Author contributions

Study Conceptualization and design: MF and design of Manish kumar, ^1,^Shweta Sahni^2^, Vivekanand^1^, Deepak kumar^3^, Neetu kushwah^4^, Divya Goel^5^ and Mohammed Faruq^3^

## STAR method

### Materials and Methods

#### Cell culture

##### Peripheral blood mononuclear cells (PBMC) isolation and LCL generation

PBMCs were isolated from one healthy (control) individual using the Histopaque-1077 (Sigma-Aldrich) gradient method according to the manufacturer’s protocol. Isolated PBMCs were then transformed to generate lymphoblastoid cell lines (LCLs) using Epstein Barr virus (EBV) following previously published protocol [10]. These LCLs were maintained in RPMI 1640 medium (Gibco) supplemented with 20% FBS (Gibco), 2 mM glutamine, and 1X Penicillin & Streptomycin (Pen-Strep) (Gibco).

##### Generation of induced pluripotent stem cells (iPSCs) from Lymphoblastoid cell line (LCL)

iPSCs generation was done utilizing Okita’s episomal plasmids, with some modifications. In brief, LCL cell lines were passaged 2 days before nucleofection and maintained in an iPSC medium containing knockout DMEM F-12 (Gibco), 20% knockout serum replacement (Gibco), 55 mM beta-mercaptoethanol (Sigma), 10 mM nonessential amino acids (Gibco), 1X glutamax (Gibco), 1X Pen-Strep, 10 ng/ml recombinant human bFGF (Gibco). On day 0, cells were counted manually, by using a hemocytometer, and nucleofection was done with 2µg of plasmid cocktail on 1 million LCLs. After which, cells were maintained in an iPSC medium containing Vitamin C (50µg/ml) + NaB (0.5Mm) on day 2. The medium was changed every alternate day. Cells were transferred to Mouse Embryonic Fibroblast (MEF) feeder cell line on the 8^th^ day in the same culture medium till the 12^th^ day. After which, sodium butyrate was withdrawn from the medium. On day 22, Human ES-like colonies were observed. Following that, iPSC colonies were passaged manually using insulin syringes through a dissection microscope, to a new MEF feeder layer dish with iPSC medium containing 10uM rock inhibitor. The iPSCs were characterized using 4 pluripotency markers. (Supplementary Figure 1). Further, Embryoid bodies (EBs) generated with these iPSCs showed all three germ layer markers (ectoderm, mesoderm, and endoderm) (Supplementary Figure. 2).

##### Neural stem cells (NSCs) generation and pan Neuronal Differentiation

Neural stem cells (NSCs) were differentiated from three Spinocerebellar ataxia type-12 iPSCs (IGIBi002-A, IGIBi003-A, and IGIBi004-A) (published previously) [11], and two Control iPSCs, ADBSi001-A [12], and one in house iPSC generated from a healthy individual (Cell line ID. 10223) (Supplementary Figure 1 and 2) by using previously published protocol [13] with slight modifications. Briefly, the iPSCs were cut manually by an insulin syringe making 100-150 clumps, then these clumps were grown in an adherent free culture dish in the presence of an iPSC medium without basic FGF. The medium was changed every alternate day, and after 5 days floating iPSCs or embryoid bodies (EBs) were observed. After the 5th day, the medium was changed to neural induction medium (DMEM/F12 (Gibco) 1X glutamax (Gibco), 1X MEM non-essential amino acids (Gibco), N2 supplement (1X) (Gibco), B27 supplement without vitamin A (2X) (Gibco), basic FGF (20ng/ml) (Gibco), EGF (10ng/ml) (Gibco), heparin (2ug/ml) (sigma), pen-strep (1X) (Gibco). After 4-5 days of culture in a neural induction medium, EBs generated rosette-like structures at their center. These structures were marked and cut manually and transferred onto laminin-coated dishes. On the 10th day, the medium was changed to neural expansion medium having knock-out DMEM/F12, 1X Stem-pro neural supplement, 1X pen-strep, 1X glutamax, FGF-2 (10ng/µl), EGF (10ng/µl). On the 12th day, the dish was covered with a homogenous population of neural stem cells (NSCs). Further, NSCs were passaged with 30-40% confluency in another laminin-coated dish (Day 0) and left overnight for cells to adhere to the surface. For pan neuron generation, on day 1, the medium was changed to neural differentiation medium containing neurobasal medium (Gibco), B27 supplement with vitamin A (2X) (Gibco), N2 supplement (1X) (Gibco), glutamax (1X) (Gibco), MEM non-essential amino acids (1X) (Gibco), penstrep (1X) (Gibco). The medium was changed every 3rd day, and mature pan neurons were observed on day 15th. Further, the neurons were characterized using mature neuronal markers TUJ1 (Beta III tubulin) and MAP2 (Supplementary Figure 3).

##### Secondary cell line maintenance

HEK293T cell line and SK-N-SH Cell line were maintained in 12% DMEM medium containing, DMEM (Gibco), 12% FBS (Fetal bovine Serum) (Gibco), 1X glutamax (Gibco), 1X penstrep (Gibco).

##### Plasmid transfection and transient overexpression

Using the manufacturer’s protocol, RAN constructs plasmids with a variable number of CAG repeats that were transfected in the HEK293T cell line using Lipofectamine TM 3000 transfection reagent (Invitrogen). In Brief, HEK293T cells were seeded in 6 well culture plates. After the cells reached 50% confluency, for each well, a mixture of Lipofectamine, P3000 reagent, and Opti-MEM (Gibco) was mixed (1.5ul + 1.5 ul + 100ul) with 1.5ug of plasmid construct. The medium was changed with fresh 10% DMEM after 6 hours of transfection.

##### RAN construct generation

To investigate if RAN translation can occur from the *PPP2R2B* gene ORF, different CAG lengths of *PPP2R2B* were amplified using SCA12 positive patient’s DNA and cloned in pcDNA 3.1 vector including three epitope tags in each reading frame at the C-terminus. Also, the plasmid was further mutated at the ATG start codon using site-directed mutagenesis. To investigate expression without the AUG start codon in expanded CAG-expressing cells.

##### Designing of novel RAN-specific antibodies & Immunofluorescence (IF) microscopy

To test if RAN proteins are getting expressed in Spinocerebellar ataxia type-12 patient-derived neural cells, we designed two polyclonal antibodies from C-terminal regions of the *PPP2R2B* gene to generate peptide for frame-specific polyclonal antibodies (Supplementary Figure 4), obtained from Biomatik. Cells were fixed with 4% paraformaldehyde (PFA) for 20 minutes followed by permeabilization with 0.1% Triton-X 100 for 10 minutes. The cells were kept in a blocking solution (1% BSA) for one hour at RT, incubated overnight at 4°C with diluted primary antibodies. The cells were washed thrice with 1X PBS (5 minutes per wash) and incubated for 1 hour with diluted secondary antibodies at RT. Counterstaining for nuclei was done with Slowfade gold reagent DAPI (Invitrogen) for 20 minutes at RT. Imaging was done using confocal microscopy (NIKON A1R) and Zoe Cell Imager microscope.

##### RNA Fluorescence in-situ hybridization (FISH)

To detect RNA/Nuclear foci inside the nucleus of Patient-derived cell lines, we used CY3-(CTG)6-CT LNA modified at nucleotide positions 2, 5, 8, 13, 16, and 19 (Takara Bio), and Stellaris RNA FISH kit (Biosearch Technologies) was used to perform RNA FISH (Fluorescence in situ hybridization) experiments. In brief, Neural Stem Cells (NSC) were grown on confocal dishes (Thermo) and then washed with 1X DPBS, fixed with fixation buffer (Stellaris RNA FISH kit) for 10 min at RT, and permeabilized with 70% ethanol for 1 hour at 4°C. Subsequently, washing was done using Wash Buffer A, and 100µl of Hybridization Buffer containing CY3-(CTG)6-CT probe (150pm/µl) was added and the confocal dish was transferred in a humidified chamber and incubated in the dark at 37°C for 16 hours. Thereafter, washing was done by Wash Buffer A, and DAPI nuclear stain (5ng/µl) was added, and incubated in the dark at 37°C for 30 min. Later, Wash Buffer B was added to the dish, and imaging was done on a confocal microscope (NIKON A1R).

##### Protein Pull Down assay

To identify different RNA binding proteins with our *PPP2R2B* specific CAG sequence, we used one expanded RAN CAG construct (CAG = 54) and one normal length RAN CAG construct (CAG= 10), as mentioned earlier. Using that template, first linear DNA was amplified using T7 promoter-specific primer and *PPP2R2B* specific Reverse primer (Supplementary Table. 1). RNA was synthesized by in vitro transcription, using HiScribe T7 RNA synthesis kit (NEB) using manufacturer’s protocol but using Biotinylated-CTP instead of dCTP, to make the resulting amplified product to bind with streptavidin. Further streptavidin was added and nuclear protein (isolated from Spinocerebellar ataxia type-12 derived NSCs and Control NSCs) was allowed to interact with RNA, and incubated at Room Temperature for 30 min, then bound proteins were isolated using the magnetic stand, and sent for Mass Spectrometry.

#### RNA seq analysis Quality control and read alignment

The initial quality check was performed using FastQC (v0.11.9), which also served to identify potential batch effects due to different sequencing sources The samples had an average of 35 million reads and a 56% GC content. FastQC’s different modules were used to analyze all files and to verify read quality, read length, GC content, N content, adapter content, and potential sources of contamination. Substandard samples, or those exhibiting adapter content, were filtered out using Cutadapt (v.3.4) [14]. The trimmed reads were aligned to the reference transcripts (Gencode Release 19 – GRCh37.p13) using the pseudo aligner Salmon (v.1.7.0) [15] with options (numGibbsSamples=30) for inferential replicates. Salmon provided quantification of each transcript in *.quant files. Post-alignment QC was performed using MultiQC (v1.11) [16] to verify the quality of alignment based on Salmon logs, FastQC reports, and Cutadapt logs.

#### Differential Expression Analysis

The *.quant files were gene length scale transformed and imported into R as a SummarizedExperiment object (v.1.24.0) [17] using tximport (v.1.22.0) [18]. The imported read counts were pre-filtered with a read-count cut-off of at least 5 reads mapped on a gene for a minimum of 3 samples. We used the DESeq2 (v.1.34) [19] R Bioconductor package for the quantification of differentially expressed genes, considering sex and batch as confounding factors. Finally, the apeglm (v1.8.0) [20]method was used to estimate a shrunken fold change for better ranking and accurate effect size of fold changes.

DESeq2 corrects batch effects based on the model design mentioned as confounding factors before performing differential expression analysis. However, the variance-stabilizing transformation function in DESeq2 does not use the model design for variance removal. Therefore, it is anticipated that variance associated with the batch or other confounding variables will persist. For data representation purposes, the transformed data was batch corrected using the removeBatchEffect function from the Limma (v3.42.2 [21] R Bioconductor package. The quality of batch correction was assessed using Principal Component Analysis (PCA).

#### Differential Transcript Expression

Salmon outputs were imported without summarizing to gene level and filtered the data for any transcripts having 10 reads for at least 3 samples. The batch effect was corrected using Limma (v3.42.2) [21] on each inferential replicate and used fishpond (2.6.0) [22] for transcript-level differential expression.

#### Image acquisition and analysis

Characterization images of iPSC, RNA-FISH, RAN translation in NSCs, and neuronal characterization images were taken on A1R confocal microscope by Nikon, and analysis was done using raw TIFF files on Fiji ImageJ software, Images of Embryoid bodies characterization, having 3 germ layer markers and HEK293T cell line images for RAN translation were taken on ZOE cell imager microscope.

#### Statistical Analysis

The statistical tests used are mentioned in the figure legends, and the results were acquired from at least three independent cell culture procedures. Student t-test was used to calculate the significance between Spinocerebellar ataxia type-12 positive and Control cell lines in PolyGlutamine and PolySerine analysis.

## Data Availability

The data used for RNA sequencing analysis and figures are available at NCBI with Bioproject PRJNA391759 and https://github.com/viv3kanand/SCA12-RNA-Seq-Analysis

